# Transgenic Inducible MHC I Overexpression in Mouse Alveolar Type 2 Cells

**DOI:** 10.1101/2025.06.11.659140

**Authors:** Justine Mathé, Sylvie Brochu, Marc K. Saba-El-Leil, Caroline Coté, Amrita Karia, Sébastien Harton, Claude Perreault

## Abstract

The major histocompatibility complex class I (MHC I) is crucial in adaptive immunity, enabling CD8+ T cells to detect and eliminate infected and cancerous cells. Recent studies have uncovered significant variability in MHC I expression across tissues, challenging the traditional belief of uniform expression. Lung epithelial cells (LECs) express meager amounts of MHC I, which preserves the lung epithelium from excessive inflammation but renders it more susceptible to cancer and infection. Despite MHC I overexpression in various immunopathologies, its precise role in disease initiation or progression remains unclear due to the absence of suitable in vivo models for studying MHC I overexpression. This study introduces a novel mouse model with targeted surface MHC I upregulation. Leveraging a conditional Cre-lox system, we augmented Nlrc5 expression to specifically upregulate MHC I in alveolar type 2 (AT2) LECs, known for their low basal expression of MHC I and significant overexpression in disease. Our model demonstrated a rapid and sustained tenfold increase in MHC I surface expression persisting for up to a year without triggering pathology or inflammation. Comprehensive characterization and validation of this model indicated that MHC I overexpression does not serve as a primary initiator of respiratory diseases under steady-state conditions and shows a therapeutic window for increasing MHC I without significant damage to the lung epithelium. This adaptable model offers insights into the effects of tissue-specific MHC I regulation and presents new avenues for therapeutic development.

## Introduction

The major histocompatibility complex class I (MHC I) is essential for adaptive immunity. By presenting peptides to CD8+ T cells, MHC I enables the rapid detection and elimination of infected or cancerous cells, ensuring continuous immune surveillance and defense (Granados et al., 2015). The MHC I complex is a heterodimeric structure composed of the MHC I heavy chain, beta-2-microglobulin (B2M), and a peptide. Its assembly is crucial for surface expression. Within cells, the MHC I-peptide complexes undergo several maturation steps. Firstly, peptides are generated by the proteasome and then transported to the endoplasmic reticulum via the TAP1 and TAP2 transporters. Subsequently, peptide loading onto the MHC I complex is facilitated by the peptide loading complex (PLC) (Blees et al., 2017). Finally, specific MHC I-peptide complexes are released on the cell surface, ready to interact with CD8+ T-cells. Transcription of MHC I is regulated by several factors, including the NOD-Like Receptor C5 (NLRC5), NF-κB, and Interferons (IFN) (Jongsma et al., 2019). Among them, NLRC5 is the principal regulator of MHC I because it upregulates MHC I per se and the transcription of critical components of the MHC I processing and peptide loading pathway (Ludigs et al., 2015).

Traditionally, MHC I was believed to be uniformly expressed by all nucleated cells at similar levels. However, recent studies have unveiled significant heterogeneity in MHC I expression across various tissues (Boegel et al., 2018). We reported that lung epithelial cells (LECs) exhibit particularly low MHC I levels, which are dramatically upregulated in chronic respiratory diseases (Mathé et al., 2022, 2024). Despite the documented overexpression of MHC I in multiple immunopathologies such as chronic respiratory diseases, inflammatory bowel diseases, neurodegenerative diseases, autoimmune myopathies, and type 1 diabetes, the precise physiological roles of MHC I in these diseases remain elusive (Allison et al., 1988; Bär et al., 2013; Cebrián et al., 2014; Fréret et al., 2013; Mathé et al., 2024). A major hurdle in studying MHC I upregulation in vivo has been the complexity of generating a suitable model. The main reasons are i) MHC I expression in the trophoblast is lethal for the fetus (Aït-Azzouzene et al., 1998; Jaffe et al., 1992; van den Elsen et al., 2001), ii) surface upregulation of MHC I through *H2-k1* knock-in is limited by the availability of B2M, peptides, and PLC genes (Allison et al., 1988; Benhammadi et al., 2020; Blees et al., 2017; Fréret et al., 2013), iii) most of the MHC I transcriptional regulators (e.g., IFN, NF-κB) have numerous functions inside the cell, outside MHC I regulation (Capece et al., 2022; Huang et al., 1993).

This study aimed to develop a mouse model with targeted surface overexpression of MHC I in a cell type with low constitutive expression, alveolar type 2 (AT2) LECs (Mathé et al., 2022). To achieve this, we engineered a mouse model where the expression of *Nlrc5* could be conditionally increased using the Cre-lox system. By crossbreeding these mice with *Sftpc-CreER* mice, in which CRE is activated in SFTPC+ cells following tamoxifen (TAM) injection, we successfully induced a substantial tenfold increase in MHC I surface expression in AT2 LECs, which are the LECs subtypes expressing the lowest amount of MHC I (Mathé et al., 2022). This elevation was detectable within five days post-injection and remained consistent for up to a year. We extensively validated the specificity and efficacy of our model and investigated the consequences of MHC I overexpression in AT2 under steady-state conditions. Our adaptable model lays the groundwork for boosting MHC I expression in diverse cell types and contexts, offering valuable insights into the upregulation of MHC I upregulation across various diseases.

## Material and methods

### Plasmid design and in vitro validation

The *LSL-Nlrc5* plasmid was constructed by VectorBuilder: vector ID VB210114-1136bna (detailed information on vectorbuilder.com). Three sets of primers were designed to validate the construct: two targeted the *Nlrc5*-*mCherry* fusion (NLRC5-CHERRY_TG-1 and 2), while the last targeted the *LSL*-*Nlrc5* fusion (LSL-NLRC5_TG-1). The sequences of these primers are the following: NLRC5-CHERRY_TG-1, “AGTGAACTTGGAGTGGAACCGGATCACAGC” (Sense) and “TGGTCTTCTTCTGCATTACGGGGCCGTCGG” (AntiSense); NLRC5-CHERRY_TG-2, “TCAGCTTGGCAGAGAATAACCTGGCTGG” (Sense) and “GAGCCGTACATGAACTGAGGGGACAGG” (AntiSense); LSL-NLRC5_TG-1, “AGCCATACCACATTTGTAGAGGTTTTACTTGC” (Sense) and “CTCCATGTCCAAAGGTACATCAAGCTCGAAGC” (AntiSense). Three plasmids containing *LSL-Nlrc5*, the Cre recombinase, and a transposase were expanded in E. coli with ampicillin selection for validation. Plasmids were then extracted using the Qiagen Midiprep Kit (Qiagen, Germantown, Maryland). Following extraction, the plasmids were transfected into highly transfectable HEK-293 cells using Lipofectamine 2000 (Invitrogen, Waltham, Massachusetts, USA). Transfections were performed with *LSL-Nlrc5* and transposase plasmids, with or without the Cre recombinase plasmid. At 48 hours post-transfection, cells were harvested, stained for 30 minutes with HLA-ABC antibodies (pan anti-human MHC I, clone W6/32), and analyzed by flow cytometry to assess mCherry and MHC I fluorescence.

### Microinjection

First, the *LSL-Nlrc5* plasmid and hyPBase (VectorBuilder VB210214-1060cbr) were amplified and purified from E.coli using a Qiagen Midiprep Kit (Qiagen, Germantown, Maryland). The DNA was redissolved in TE buffer (pH 7.4) in equal molar ratios, and the final concentration of both plasmids was 25 ng/µL. Transgenic mice were produced by microinjecting the purified plasmids (1-2 ng/ul) into the pronucleus of zygotes (0.5 days post coitum) and were transferred to pseudopregnant females by surgical transfer the next day. The embryos developed to term, and resulting pups were genotyped by genomic PCR using three primer sets to identify the “founder” mice. After several selection steps (described below), the *LSL-Nlrc5* colony was generated by crossing the selected founder with C57Bl/6 mice.

### Insertion quantification

We performed qPCR using a non-intron-spanning strategy to evaluate the number of transgene copies in each *LSL-Nlrc5* positive mouse. DNA was extracted from mouse ear tissue using REDExtract-N-Amp tissue PCR (Sigma-Aldrich, St. Louis, Missouri). DNA was amplified using assays designed with the Universal Probe Library (UPL) from Roche (www.universalprobelibrary.com). For each qPCR assay, a standard curve was performed to ensure that the efficiency of the assay was between 90% and 110%. Primer sequences are as follows: *Nlrc5* : Forward: AAGCTCCTGAAGACCTCTGC and reverse: GAGTAAGCCATGCTCCTGGG) used with UPL probe#59, *Tert* (Forward: GCTCACAGCCTCCACCTG and reverse: CGCACTATCCCACGTCCT). The QuantStudio qPCR instrument (Thermo Scientific, Waltham, Massachusetts) was used to detect the amplification level. The number of copies was determined using *Tert* as a reference gene.

### Mice

C57BL/6J (B6) and B6.129S-Sftpctm1(cre/ERT2)Blh/J (*Sftpc-CreER*) mice were purchased from The Jackson Laboratory (Bar Harbor, Maine). The Texas A&M Health Sciences provided *Nlrc5* KO mice generated at the Dana-Farber Cancer Institute (Biswas et al., 2012). *Sftpc-CreER:LSL-Nlrc5* (TG) mice were obtained by crossing *LSL-Nlrc5* mice with *Sftpc-CreER* mice. WT littermates were used as controls. All mice descending from *Sftpc-CreER* and *LSL-Nlrc5* mice were genotyped at two weeks old by genomic PCR using the three sets of primers described above. Twenty-one-day-old TG and WT mice received one intraperitoneal injection per day of 1 mg TAM diluted in 50 µL corn oil for five days. Mice’s weight was monitored daily during the injection period, weekly for the following week, and monthly for mice that grew older. A pilot study evaluated the toxicity of the injection to determine the appropriate dosage, with mortality rates varying between 0 and 20%. Mice aged five weeks to 1 year were analyzed. Unless otherwise specified, each analysis included males and females in similar proportions. Mice were maintained under specific pathogen-free conditions at the Institute for Research in Immunology and Cancer. All mice were cared for and handled according to the Canadian Council on Animal Care guidelines. All protocols were approved by the Comité de Déontologie de l’Expérimentation sur les Animaux de l’Université de Montréal.

### Lung digestion

Lung cells were obtained, as previously reported (Benhammadi et al., 2020; Mathé et al., 2022). Briefly, lung tissue was minced using scissors in RPMI-1640 medium (Invitrogen, Waltham, Massachusetts). The tissue fragments were then enzymatically digested with 0.01% Liberase Thermolysin Medium (Roche Applied Science, Penzberg, Germany) and 0.1% DNase I (Sigma-Aldrich, St. Louis, Missouri) in RPMI-1640 (Invitrogen, Waltham, Massachusetts). The resulting cell suspensions were maintained at 4°C in FACS buffer (PBS, 0.1% (w/v) BSA, and 0.02% (w/v) NaN3) before cell staining for flow cytometric analysis and cell sorting.

### Flow cytometry and cell sorting

Cells obtained from enzymatic digestion were stained with specific antibodies. For LECs, the following antibodies were used: anti-Epcam APC-Cy7 (Biolegend, San Diego, California, cat #118218), anti-CD45 APC (Biolegend, cat #559864), anti-IA/IE Alexa Fluor 700 (clone M5/114.15.2, Biolegend, San Diego, California, cat #107622), anti-CD31 Alexa Fluor 488 (Biolegend, cat #102414), pro-SPC (Fisher Scientific, Waltham, Massachusetts, cat #AB3786MI). We used anti-CD45 eFluor 605NC (eBiosciences, cat #93045173), anti-CD3 Pacific Blue (Biolegend, San Diego, California, cat #100213), and anti-CD8 APC-CY7 (Biolegend, San Diego, California, cat #100713) for leukocyte staining. Anti-H2-K^b^/D^b^-PE (clone 28-8-6; BioLegend, San Diego, California, cat #553575) was used to stain for MHC I. Cells stained with isotype control antibodies (BioLegend) helped exclude non-specific binding. Viability was determined using a live/dead staining method. Flow cytometry analysis and cell sorting were conducted using a ZE5 Cell Analyzer (Bio-Rad Laboratories, Mississauga, Ontario, Canada) and a FACSAria (BD Biosciences, Franklin Lakes, New Jersey). Data analysis was performed with FlowJo software V9.

### Immunohistochemistry

Lungs were fixed in 10% formalin, embedded in paraffin, and sectioned into 4-μm slices. The sections were stained with H&E and scanned using the NanoZoomer Digital Pathology system (Hamamatsu, Shizuoka, Japan) at 40X magnification. Image analysis was performed with NDP.view software (version 2.7.52, Hamamatsu).

### Quantitative RT-PCR analysis

RNA integrity was validated using a Bioanalyzer 2100 (Agilent Technologies, Santa Clara, California). Total RNA was treated with DNase and reverse transcribed using the Maxima First Strand cDNA synthesis kit with ds DNase (Thermo Scientific, Waltham, Massachusetts). Gene expression was determined using assays designed with the Universal Probe Library from Roche (Roche, Penzberg, Germany). For each qPCR assay, a standard curve was performed to ensure that the efficiency of the assay was between 90% and 110%. Primer sequences are as follow: *Psmb9* (ref: NM_013585) forward (gcccaagccatagctgac) and reverse (gatgttcttcaccacgtttgc) used with UPL probe#78, *H2-d1* (ref: NM_010380.3) forward (ggtggaaaaggaggggacta) and reverse (gcagctgtcttcacgcttt) used with UPL probe#106, *Tap1* (ref: NM_013683) forward (ttccctcagggctatgacac) and reverse (ctgtcgctgacctcctgac) used with UPL probe#48, *Psmb8* (ref: NM_010724.2) forward (cacactcgccttcaagttcc) and reverse (tctcgatcactttgttcatcctt) used with UPL probe#81, *Tap2* (ref: NM_011530) forward (ggtggcctgctctccttc) and reverse (ccgtacatgtaaaccaggttcc) used with UPL probe#69, *Psmb10* (ref: NM_013640.3) forward (gttccagccaaacatgacg) and reverse (cagagcccaggtcactcag) used with UPL probe#31, Nlrc5 (ref: NM_001033207.3) forward (gtactcacatttgcccaggag) and reverse (gacacgaagctgataagtggtg) used with UPL probe#15, *Stat1* (ref: NM_001205314.1 and NM_001205313.1) forward (agctcgtggagtggaagc) and reverse (ggtctctgcaacaatggtga) used with UPL probe#74, *B2m* (ref: NM_009283.4) forward (CAGCAAGGACTGGTCTTTCTAT) and reverse (AACTCTGCAGGCGTATGTATC), *H2-k1* (ref: NM_001001892.1 and NM_001001892.2) forward (ATCACCCTGACCTGGCAGT) and reverse (CCACTTCTGGAAGGTTCCATC) used with UPL probe#54, *Gapdh* (ref: NM_008084.2) forward (tgtccgtcgtggatctgac) and reverse (cctgcttcaccaccttcttg) used with UPL probe#80, *Gusb* (ref: NM_010368.1) forward (aaaatcaccctgcggttgt) and reverse (tgtgggtgatcagcgtctt) used with UPL probe#6. The QuantStudio qPCR instrument (Thermo Scientific, Waltham, Massachusetts) detected the amplification level. Relative expression comparison (RQ = 2^-ΔΔCT^) was calculated using the Expression Suite software (Thermo Scientific, Waltham, Massachusetts), using *Gapdh* and *Gusb* as endogenous controls. Data were normalized to WT mice of the same sex. Unless otherwise indicated, analyses were conducted on 5-week-old mice.

### RNA-sequencing

RNA-sequencing experiments assessing *Nlrc5* and *Sftpc* expression were conducted as outlined in previous studies (Benhammadi et al., 2020; Mathé et al., 2022). Detailed methodologies are provided in the respective original publications. Raw data can be accessed on the Gene Expression Omnibus under the accession numbers GSE144722, which pertains to the comparison among thymic, colon, skin, and LECs, and GSE176228, which focuses on LECs comparison. Transcriptomic data from mice tissues were obtained from the Tabula Muris project (Schaum et al., 2018).

### Statistical analysis

Unless specified otherwise in the figure legends, data are presented as mean ± SD, with a minimum sample size of n = 3 for statistical analyses. We employed a two-tailed Student t-test using GraphPad Prism version 8 software to assess statistical differences between the two groups. For analyses involving three or more groups, we performed repeated measures of one-way ANOVA, followed by a post hoc test (Tukey or Sidak) for multiple comparisons. Statistical significance was considered at p-values ≤ 0.05 (*), ≤ 0.01 (**), ≤ 0.001 (***), ≤ 0.0001 (****), and nonsignificant results were denoted as (ns). Details regarding sample size, experimental replicates, and other pertinent information can be found in the figure legends.

## Results

### Characterization of *Nlrc5* and *Sftpc* Expression in the Lungs of B6 Mice

This study aimed to generate a mouse model overexpressing surface MHC I in AT2 LECs. We selected NLRC5 as the top candidate to facilitate MHC I upregulation. This choice was based on NLRC5’s ability to specifically increase MHC I without affecting MHC II and to enhance the expression of genes crucial for MHC I surface expression (e.g., *B2m*, *Tap1*, *Psmb9*) (Ludigs et al., 2015). Most research on NLRC5 has focused on hematolymphoid cells, leaving its role poorly investigated in other cell types (Biswas et al., 2012; Ludigs et al., 2015; Meissner et al., 2010). This prompted us first to evaluate NLRC5 expression in the lung and its role in MHC I regulation in LECs from mice. LECs were sorted using the expression levels of EpCAM and MHC class II (IA/IE) as validated in previous work (Hasegawa et al., 2017; Mathé et al., 2022). Three types of LECs were identified: Alveolar type 1 (AT1, EpCAM^int^ IA/IE^lo/neg^), AT2 (EpCAM^int^ IA/IE^hi^), and bronchiolar ECs (Bronc., EpCAM^hi^ IA/IE^lo/neg^) (Fig. 1A). The three LEC-types analyzed expressed *Nlrc5* transcripts (Fig. 1B). Notably, *Nlrc5* expression in LECs is low compared to medullary ECs from the thymus (mTECs), which express high levels of MHC I (Benhammadi et al., 2020). Flow cytometry experiments on LECs from *Nlrc5* knock-out (KO) showed a significant decrease in MHC I surface expression in *Nlrc5* KO mice compared to WT mice (Fig. 1C). These data confirmed that NLRC5 regulates MHC I expression in mouse LECs, making it a suitable model for our transgenic mice.

**Figure 1.**
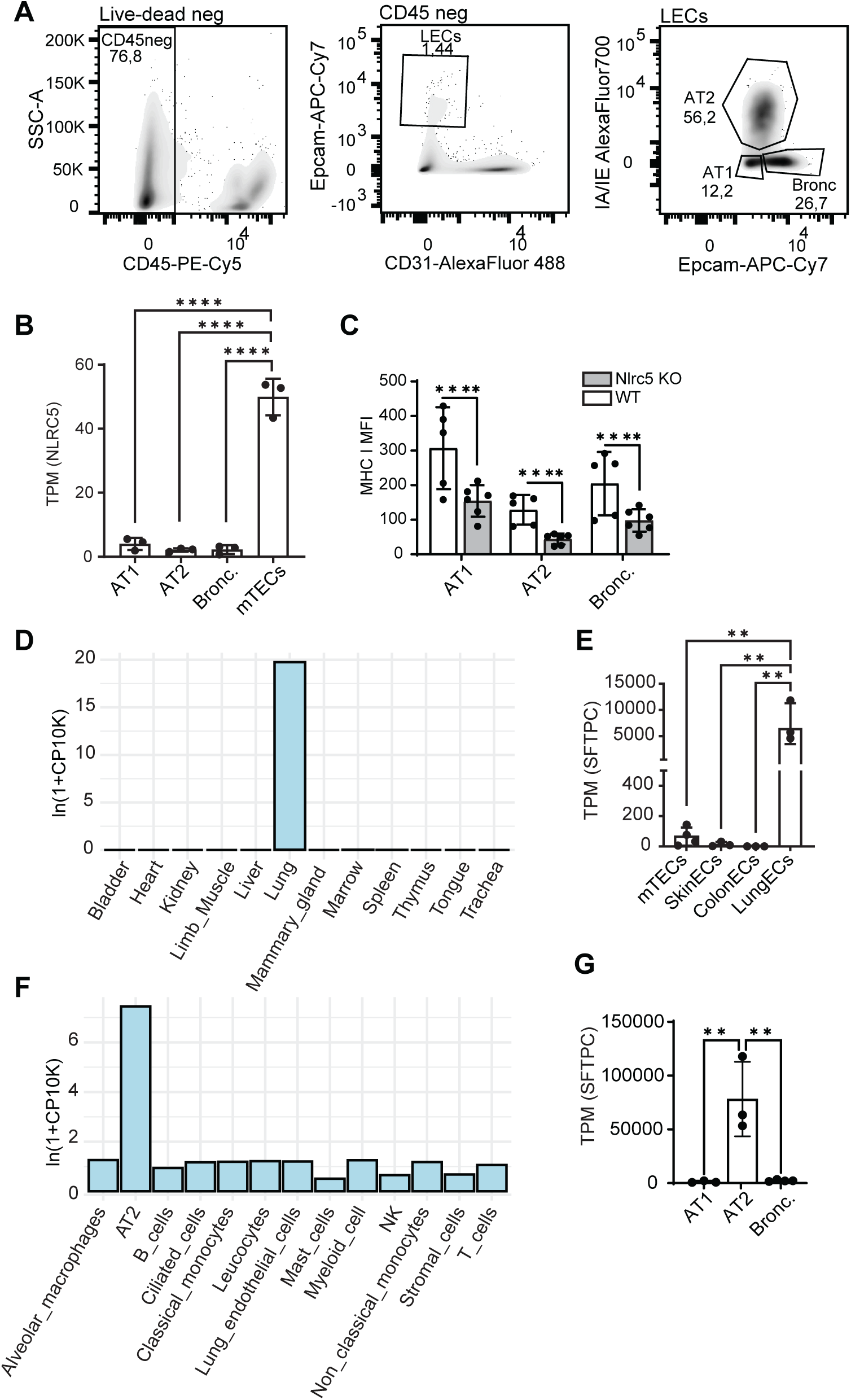
Expression of *Nlrc5* and *Sftpc* in mice. A) Gating strategy for AT1 (IA/IE^lo^ EpCAM^lo^), AT2 (IA/IE^hi^ EpCAM^int^), and bronchiolar ECs (Bronc, IA/IE^lo^ EpCAM^hi^) analyses and sorting by FACS in mice. CD31^+^ and CD45^+^ cells (endothelial cells and leukocytes) were excluded to prevent contamination of LEC populations. B) *Nlrc5* mRNA expression obtained by RNA-seq from isolated LECs and mTECs (GSE144722 and GSE176228, n=3). C) Mean fluorescence intensity (MFI) of MHC I (H2-K^b^/D^b^) in WT B6 mice (white) compared to *Nlrc5* KO mice (grey) obtained by flow cytometry (n=6). D and F) *Sftpc* mRNA expression in different mice tissues (D) and lung cells (F) from *Tabula Muris* (Schaum et al., 2018). E and G) *Sftpc* mRNA expression obtained by RNA sequencing in different mice tissues (E) and LECs (G) (GSE144722 and GSE176228, n=3). For figures A-C, E, and G, 8-week-old males were analyzed. Statistical significance was evaluated using a two-tailed Student’s t-test. p-values ≤ 0.05 (*), ≤ 0.01 (**), ≤ 0.001 (***), and ≤ 0.0001 (****) are indicated. TPM, transcripts per million.

Then, we aimed to select a promoter specifically expressed in LECs to increase NLRC5 expression in these cells. Many genes are used in the literature as markers for different LEC types. For example, AQP5 identifies AT1 cells but is also expressed in other organ cell types (Wittekindt & Dietl, 2019). *SFTPC*, which codes for surfactant protein C, is a widely used marker of AT2 cells (Nureki et al., 2018). AT2 is the LEC type presenting the lowest amount of MHC I. It represents 60% of LECs from the alveoli, making it an interesting choice to investigate the impact of MHC I upregulation on lung epithelium (Wang et al., 2007). To confirm the specificity of *Sftpc* to AT2 cells, we examined its expression in several mice tissues from the *Tabula Muris* project (Schaum et al., 2018). Our analysis showed that *Sftpc* is exclusively expressed in the lung (Fig. 1D). We examined its expression in RNA-seq data from several ECs obtained in our laboratory (thymus, skin, colon, and lung ECs). Our analysis revealed specific and high expression of *Sftpc* in LECs, with almost no expression in ECs from other tissues (Fig. 1E). Only mTECs exhibited low but detectable expression of *Sftpc* mRNAs, which could be explained by the promiscuous gene expression in these cells that facilitates tolerance establishment (St-Pierre et al., 2015). By exploring *Tabula Muris* data, we observed that in the lung, AT2 is the only cell type expressing a substantial amount of *Sftpc* (Fig. 1F). RNA-seq data analysis from sorted LECs confirmed the specificity of *Sftpc* to AT2 cells, showing almost undetectable levels of *Sftpc* mRNAs in AT1 and bronc—ECs (Fig. 1G).

Therefore, we concluded that NLRC5 can regulate MHC I surface expression in LECs and that the *Sftpc* is specific to AT2 cells and highly expressed in these cells. We, therefore, inferred that the overexpression of *Nlrc5* under the control of the *Sftpc* promoter should effectively increase MHC I levels in AT2 cells in vivo.

### Generation of *LSL-Nlrc5* mice

To create our transgenic mouse model overexpressing NLRC5 in AT2 cells, we designed a plasmid containing the *Nlrc5* gene fused to mCherry (Fig. 2A). The mCherry tag allows us to detect the transgene and distinguish transgenic NLRC5 from endogenous NLRC5. Since the functional domain of NLRC5 is at the N-terminal end, we inserted mCherry in the C-terminal to avoid altering its function.

**Figure 2.**
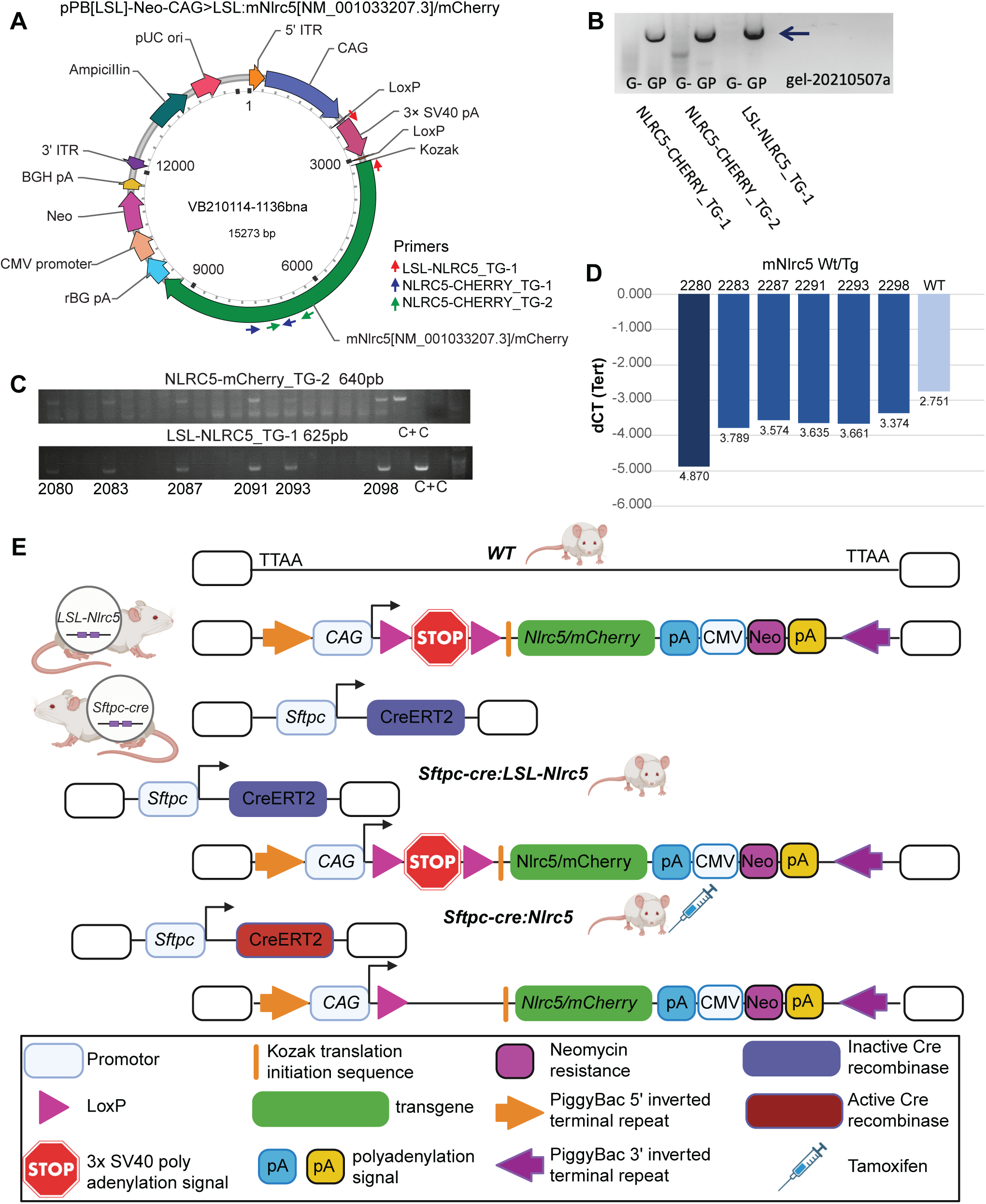
Generation of *LSL-Nlrc5* mice. A) Schematic representation of the vector construction provided by VectorBuilder. B) PCR vector validation using three sets of primers (NLRC5-CHERRY_TG-1, NLRC5-CHERRY_TG-2, and LSL-NLRC5_TG-1). Primer’s localization is indicated by arrows on panel A. Genomic DNA (G) or genomic DNA plus plasmid (GP) was used to determine the specificity of the primer toward transgenic *Nlrc5*. C) Genotyping of mice obtained following microinjections of the vector in B6 embryos. The sex of the *LSL-Nlrc5* mice is depicted in Table S1. D) ΔCT (cycle threshold) of *Nlrc5*, obtained by qPCR, in B6 (WT) mice and mice carrying the *LSL-Nlrc5* sequence after the microinjections. E) The structure of targeted alleles in WT, *LSL-Nlrc5*, *Sftpc-creER*, *Sftpc-creER:LSL-Nlrc5,* and *Sftpc-creER:Nlrc5* (treated with TAM) mice. The transgene was inserted using a PiggyBAC vector into TTAA sequences of the genome.

To induce NLRC5 upregulation in AT2, we used *Sftpc* as a promoter. Studies reported that MHC I expression in the trophoblast induces abortion of the fetus (Aït-Azzouzene et al., 1998; van den Elsen et al., 2001). SFTPC is known to be expressed in placenta tissues (Salminen et al., 2009; Sati et al., 2010). To avoid potential lethality of the fetus during gestation, we add a Lox stop Lox (LSL) sequence before the *Nlrc5-mCherry* gene to repress its constitutive expression. In cells with active Cre recombinase, the LSL sequence will be cleaved and NLRC5-mCherry expressed. We inserted our construct into a PiggyBac vector, enabling efficient and specific genomic transposition while minimizing the risk of mutagenesis (Kim & Pyykko, 2011). We added neomycin and ampicillin resistance genes for in vitro validation of the model. We designed three primers to validate our plasmid (Fig. 2A-B). After confirming the induction of NLRC5-mCherry in vitro following plasmid and Cre transfection (Fig. S1), we performed microinjections to obtain mice carrying the *LSL-Nlrc5* transgene. Six *LSL-Nlrc5* positive mice were obtained following microinjections (Fig. 2C). These mice underwent several validation steps, including assessments of MHC I expression levels, the proportion of AT2 cells expressing the transgene, and transmission to progeny, as described below.

First, the six potential founders carried one or more insertions of our transgene. We used non-intron-spanning qPCR to evaluate the number of transgene copies in each mouse (Fig. 2D). By comparing the cycle threshold (CT) between *LSL-Nlrc5* founders (unknown number of *Nlrc5* copies) and B6 mice (two endogenous copies of *Nlrc5*) mice, we were able to evaluate the number of copies of *Nlrc5* in each *LSL-Nlrc5* founder. All mice, except 2280, had one transgene insertion (two endogenous copies of *Nlrc5* plus one transgenic copy). We selected only the mice carrying one insertion to ensure a homogeneous distribution of the transgene in the progeny. Selected mice were bred with *Sftpc-CreER* mice (which express TAM-inducible Cre in Sftpc+ cells) to produce *Sftpc-CreER:LSL-Nlrc5* (TG) mice (Fig. 2E).

### Selection of the Optimal Founder

To select the best potential founder to generate the LSL-NLRC5 colony, we analyzed several parameters among the five mice carrying one insertion of our transgene. First, we evaluated the transmission rate of the transgene to the progeny. According to Mendelian law, a mouse with one transgene insertion should produce 50% transgenic offspring. Three of the five pre-selected mice (2283, 2287, 2298) transmitted the transgene to approximately 50% of their progeny (Table S1). These mice were selected for further analysis. Additionally, we measured the weight of TG and WT mice during and after TAM injection to ensure that the transgene expression did not induce systemic defect. TG and WT mice showed no weight differences while receiving TAM treatment (Fig. 3A).

**Figure 3.**
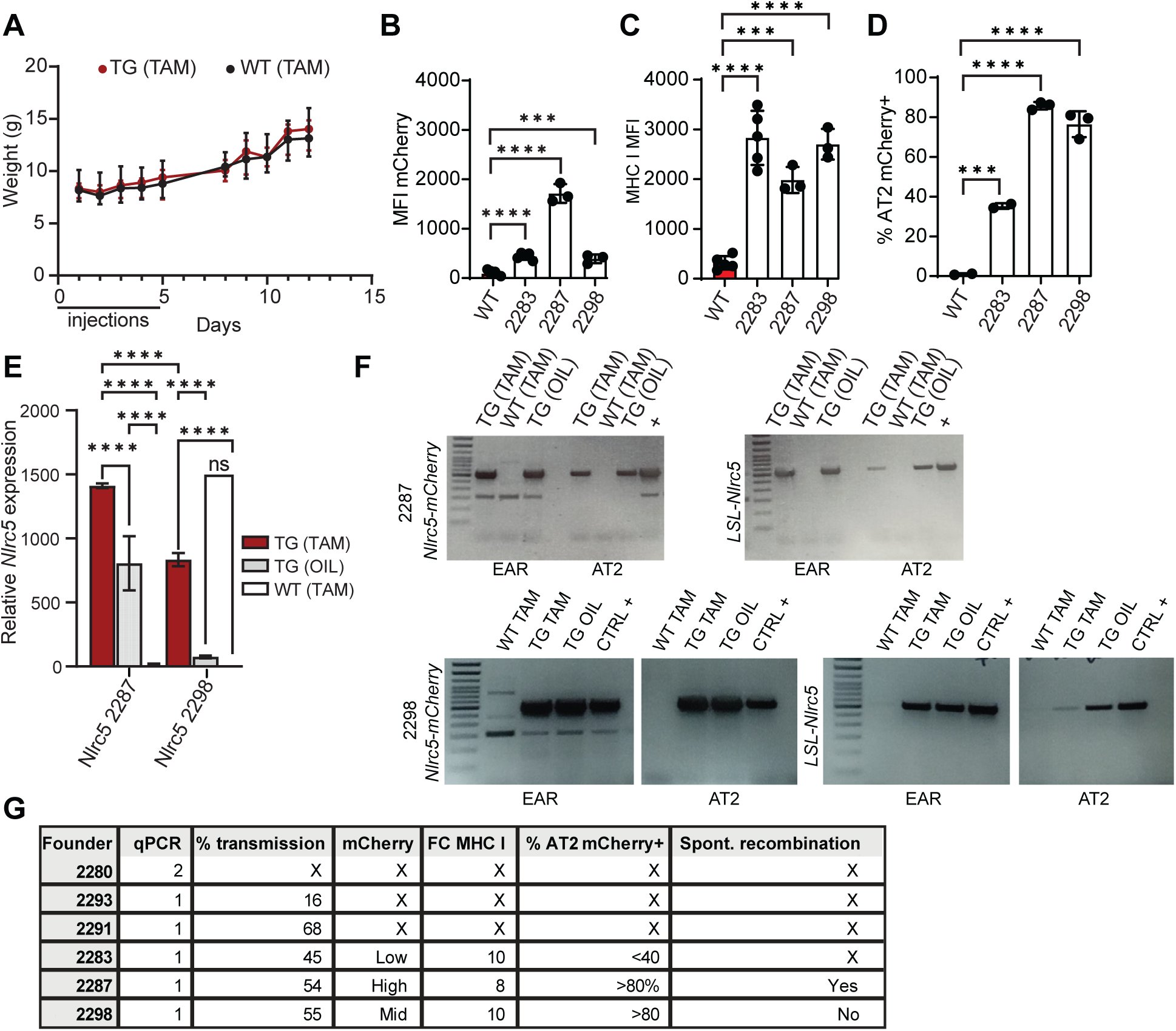
Validation of the model and selection of the founder. A) Weight of TG and WT littermate mice during the TAM injections period and the following seven days (n = 5). B-C) Mean fluorescence intensity (MFI) of mCherry (B) and MHC I (C) in AT2 from WT and TG mice treated with TAM. D) Percentage of AT2 expressing mCherry in TG and WT mice treated with TAM. Descendants from the founders 2283, 2287, and 2298 were analyzed (n=3). E) Relative mRNA expression of *Nlrc5* in AT2 from TG or WT mice treated with TAM or oil obtained by quantitative RT-PCR (n=3). Statistical significance was evaluated using a two-tailed Student’s t-test. p-values ≤ 0.05 (*), ≤ 0.01 (**), ≤ 0.001 (***), ≤ 0.0001 (****). F) PCR genotyping was performed on sorted AT2 or ear DNA from TG or WT mice treated with TAM or oil. DNA was extracted from the same number of AT2 cells for each group. G) Summary of all the criteria that led to selecting one founder for the rest of the study.

A critical criterion was the level of transgene expression and MHC I in TG mice treated with TAM. Flow cytometry analysis showed that TG descendants from 2283, 2287, and 2298 expressed mCherry and significantly increased MHC I levels following TAM treatment (Fig. 3B-C). Interestingly, the two mice with the lowest mCherry expression (2283 and 2298) exhibited the highest MHC I level. Then, we evaluated the percentage of AT2 cells expressing the transgene in these mice. Flow cytometry analysis revealed that descendants from 2287 and 2298 showed mCherry expression in more than 80% of their AT2 cells (Fig. 1D). In contrast, descendants from 2283 had an expression rate of less than 40%. Thus, 2283 was not considered the best choice for a founder. To determine whether 2287 or 2298 was more suitable as founder, we evaluated the *Nlrc5* increase induced by TAM injection using qPCR (Fig.3E). We also assessed the spontaneous recombination rate by measuring *Nlrc5* expression in transgenic mice that received only oil (TAM vehicle) injections. Previous studies reported up to 25% spontaneous recombination in *Sftpc-CreER* mice (Rock et al., 2011). Our data showed significant upregulation of MHC I following TAM injection in mice from both founders (Fig.3E). 2287 showed significant spontaneous recombination, as indicated by increased *Nlrc5* expression in mice injected with oil. This was not observed in 2298, which appeared as the best choice as a founder. We performed genomic PCR on sorted AT2 or ear DNA from 2298 TG descendant mice treated with TAM or oil and WT mice treated with TAM. Our data confirmed the cleavage of the LSL sequence only in AT2 from *LSL-Nlrc5* mice treated with TAM (Fig. 3F). Therefore, mouse 2298, which met all the criteria for a good founder, was selected and used to generate the *LSL-Nlrc5* colony used for subsequent analyses (Fig. 3G).

### MHC I overexpression is stable and specific to AT2 in TG mice

Next, we investigated the specificity of transgene induction in the progeny of 2298 crossed with *Sftpc-CreER* mice. Theoretically, mCherry fluorescence should be specific to AT2 cells following TAM treatment in these mice.

We performed flow cytometry analysis on LECs, evaluating the percentage of mCherry+ MHC I^hi^ cells in WT and TG mice treated with TAM. Only AT2 cells showed a significant increase in mCherry+ MHC I^hi^ cells in TG mice treated with TAM compared to WT mice (Fig. 4A). A small proportion of AT1 cells (4.22%) and bronchiolar ECs (2.78%) also expressed the transgene (*mCherry+*), likely due to a minor population of these cells expressing *Sftpc* (Figure 1F-G). Given that AT2 cells can differentiate into AT1 and specific bronchiolar ECs subtypes, a small percentage of these cells were expected to express the transgene (Barkauskas et al., 2013; Jansing et al., 2017). Finally, we examined mCherry expression outside the LEC population and found it to be nearly absent in CD45+ cells (leukocytes; 0.94% expressing mCherry) and CD31+ cells (endothelial cells; 0.18% expressing mCherry) (Fig. 4B). Therefore, NLRC5-mCherry expression is specific to AT2 cells in our TG mice treated with TAM.

**Figure 4.**
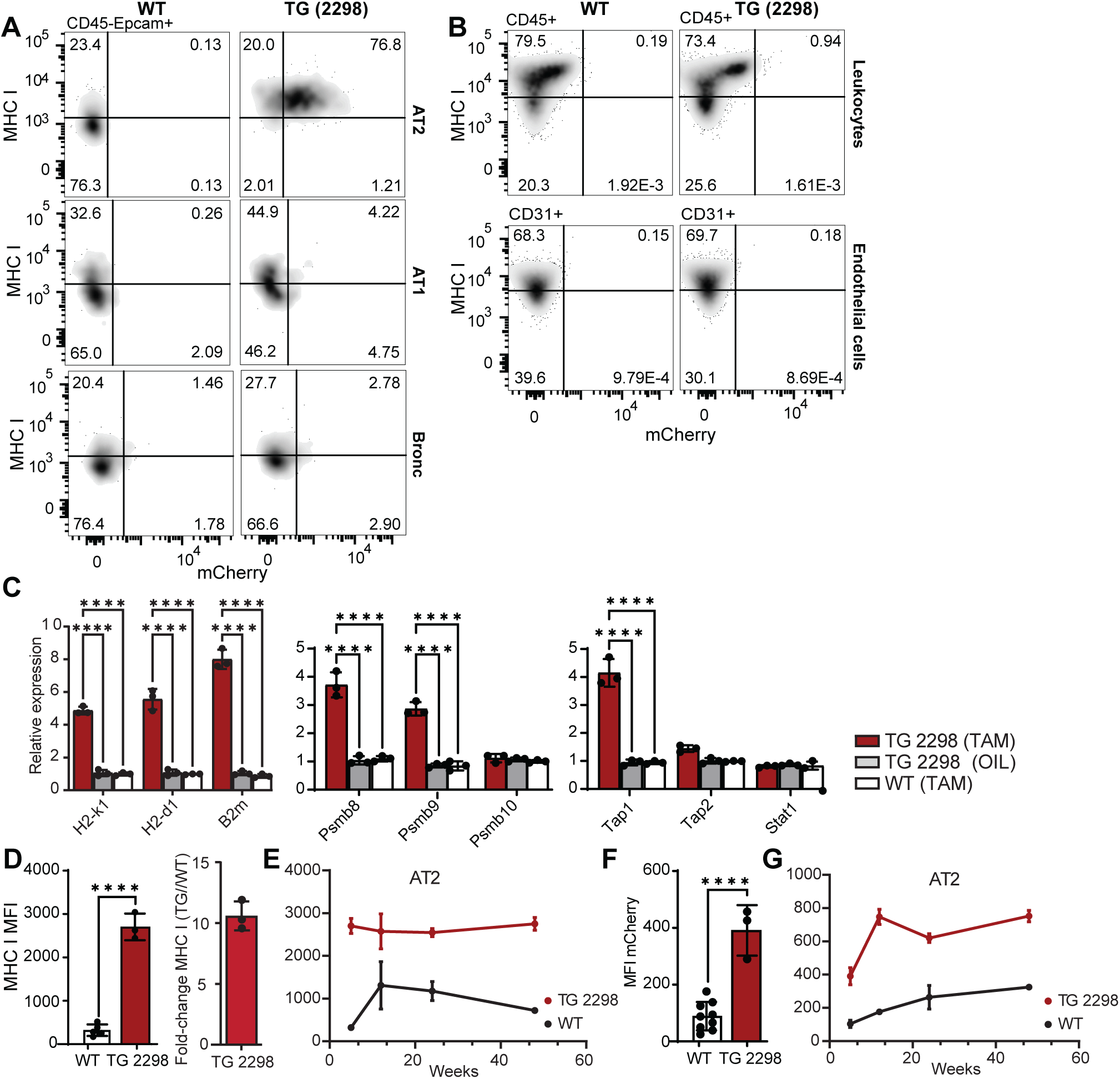
MHC I upregulation is stable and specific to AT2 in TAM-induced TG mice. A) Percentage of mCherry^+^ MHC I^hi^ LECs (AT1, AT2, and Bronchiolar ECs (Bronc.)) in the lungs of TG and WT TAM-treated mice evaluated by flow cytometry. B) Percentage of CD45+ (leukocytes) and CD31+ (endothelial) cells expressing mCherry and MHC I in WT and TG mice treated with TAM. C) Relative mRNA expression obtained by RT-qPCR of MHC I genes (*H2-k1*, *H2-d1*, *B2m*), proteasome subunits (*Psmb8*, *9*, *10*), and PLC genes (*Tap1*, *Tap2*, and *Stat1*) in WT or TG mice treated with TAM or oil (n=3). E) Mean fluorescence intensity (MFI) (left panel) and fold-change of MHC I expression (right panel) from 5-week-old WT and TG mice treated with TAM. F) Mean fluorescence intensity (MFI) of MHC I from WT and TG mice treated with TAM at different ages (5, 8, 12, 24 and 48 weeks). G-H) Mean fluorescence intensity (MFI) of mCherry in AT2 cells from WT and TG mice treated with TAM at 5 weeks old (left panel) and at different ages (5, 8, 12, 24 and 48 weeks) (right panel) (n=3). Statistical significance was evaluated using a two-tailed Student’s t-test. p-values ≤ 0.05 (*), ≤ 0.01 (**), ≤ 0.001 (***), ≤ 0.0001 (****).

To better characterize our model, we evaluate expression levels of *H2-k1* and H2-*d1* (MHC I), *Psmb8*, *Psmb9*, and *Tap1* transcripts in TAM- or oil-treated TG mice. These transcripts are crucial for the peptide loading of MHC I and are known to be upregulated by NLRC5. Our results confirmed the upregulation of these genes in transgenic mice, demonstrating that NLRC5 not only increases MHC I in AT2 cells but also induces the transcriptional upregulation of *Psmb8*, *Psmb9*, and *Tap1* (Fig. 4C). This enhanced the expression of MHC I on the cell surface, as shown in our flow cytometry experiments (Fig. 3C and 4D).

A critical point was to evaluate the long-term impact of MHC I upregulation on lung homeostasis. Therefore, we assessed MHC I and mCherry expression in TG and WT mice treated with TAM at different ages. Our data show that MHC I overexpression was induced as early as one-week post-injection (5 weeks old), reaching a fold increase of approximately tenfold (Fig. 4D Similar results were observed for mCherry expression (Fig.4F). Moreover, we showed that NLRC5-mCherry and MHC I overexpression were stable for at least a year in steady-state conditions (Fig. 4E-G).

Overall, we have created a transgenic mouse model that overexpresses MHC I on the surface of AT2 cells following TAM treatment. This increase is specific to AT2 cells and remains stable for at least one year following the injections.

### Impact of MHC I Overexpression on LECs Homeostasis under Steady States Conditions

Theoretically, increased MHC I surface expression in AT2 cells could impact LEC homeostasis in several ways. It could heighten their susceptibility to apoptosis by making them more vulnerable to CD8+ cytotoxic T-cells (Cebrián et al., 2014). This apoptosis might also affect AT1 and Bronc ECs because of the capacity of AT2 cells to differentiate into these two cell types. We evaluated the proportion of the three LEC types in our TG mice compared to WT mice treated with TAM over time. Interestingly, the percentages of each LEC population did not significantly differ between WT and TG mice under steady-state conditions (Fig. 5A). WT and TG mice tended to increase the percentages of AT1 and Bronc. ECs over time, while AT2 cells decreased. Next, we performed Annexin-V staining to evaluate cell death within the LEC populations from the same mice (Fig. 5B-C). Our results revealed no differences between WT and TG mice over a year under steady-state conditions, indicating that the expected increase in apoptosis did not occur.

**Figure 5.**
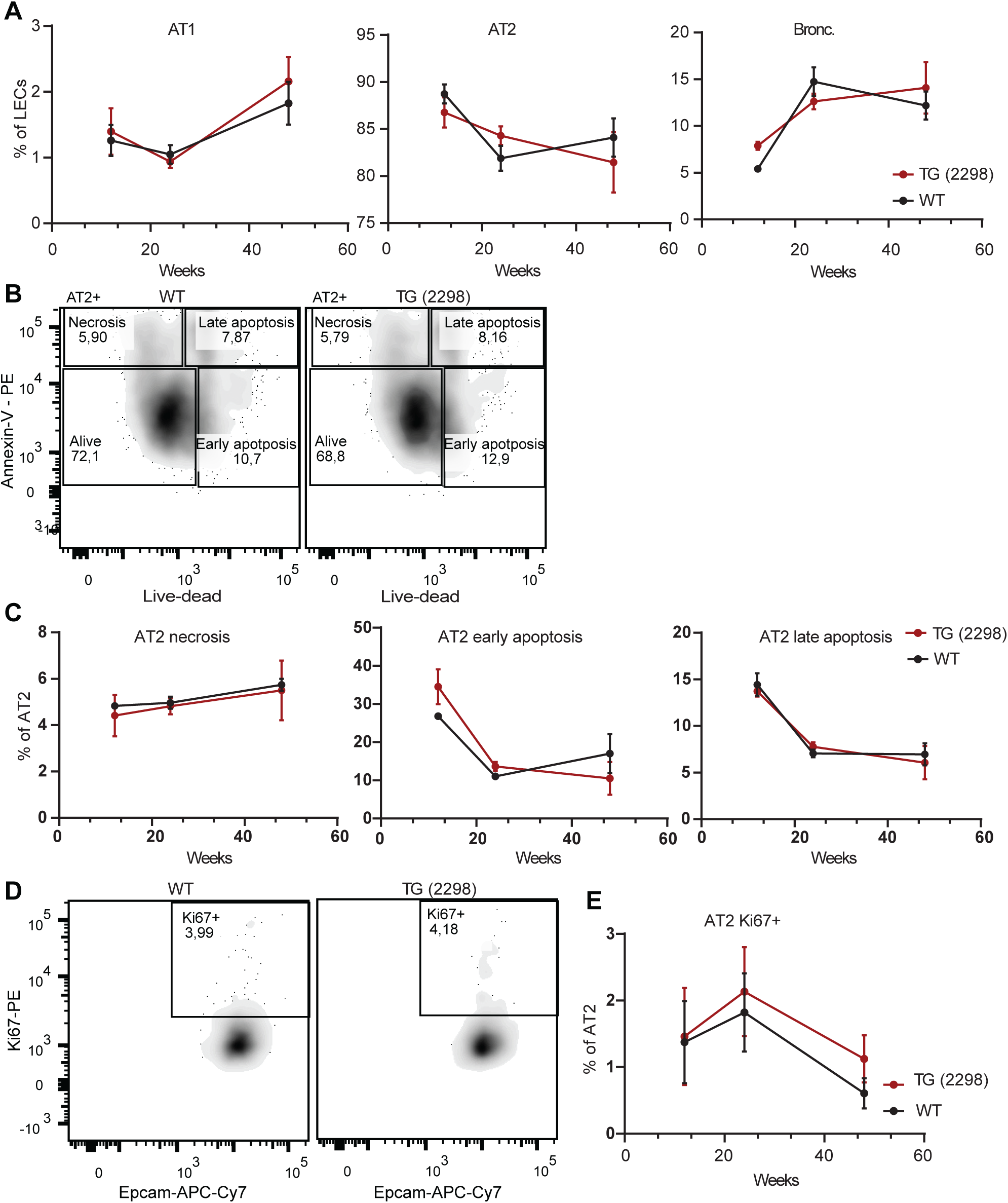
LEC fates are unaffected in TAM-treated TG mice. A) Percentage of AT1, AT2, and Bronchiolar ECs (Bronc.). B) Annexin-V staining gating strategy to assess AT2 apoptosis and necrosis in 24-week-old WT and TG mice. C) Percentage of necrotic (AnnexinV+Live-dead-), early (AnnexinV-Live-dead+), and late apoptotic (AnnexinV+Live-dead+) AT2. D-E) Gating strategy (D) and percentage (E) of proliferating AT2 Ki67+. All data were obtained from WT and TG TAM-treated mice (n=3).

When LECs are injured, AT2 cells rapidly proliferate and differentiate to replace the injured cells, particularly alveolar ECs and some bronchiolar ECs. Therefore, we performed Ki67 staining to evaluate the percentage of proliferating AT2 cells in our TG mice compared to WT mice after TAM treatment. The proportion of proliferating AT2 cells appeared slightly higher in TG mice than in WT mice, though the differences were not statistically significant (Fig. 5D-E).

Given MHC I’s role in presenting peptides and activating CD8+ T-cells, we also investigated whether the increase in MHC I led to a heightened proportion of T-cells in the lung. We evaluated the percentage of CD3+ and CD3+CD8+ T-cells in the lungs of TG and WT mice treated with TAM at different ages using flow cytometry. Our data demonstrated that TG mice did not exhibit higher amounts of CD3+ and CD8+ T-cells over one year under steady-state conditions (Fig. 6A). To characterize potential inflammation further, we performed hematoxylin and eosin staining on the lungs of TG and WT mice treated with TAM. At six months, TG mice showed no significant differences from WT mice (Fig. 6B).

**Figure 6.**
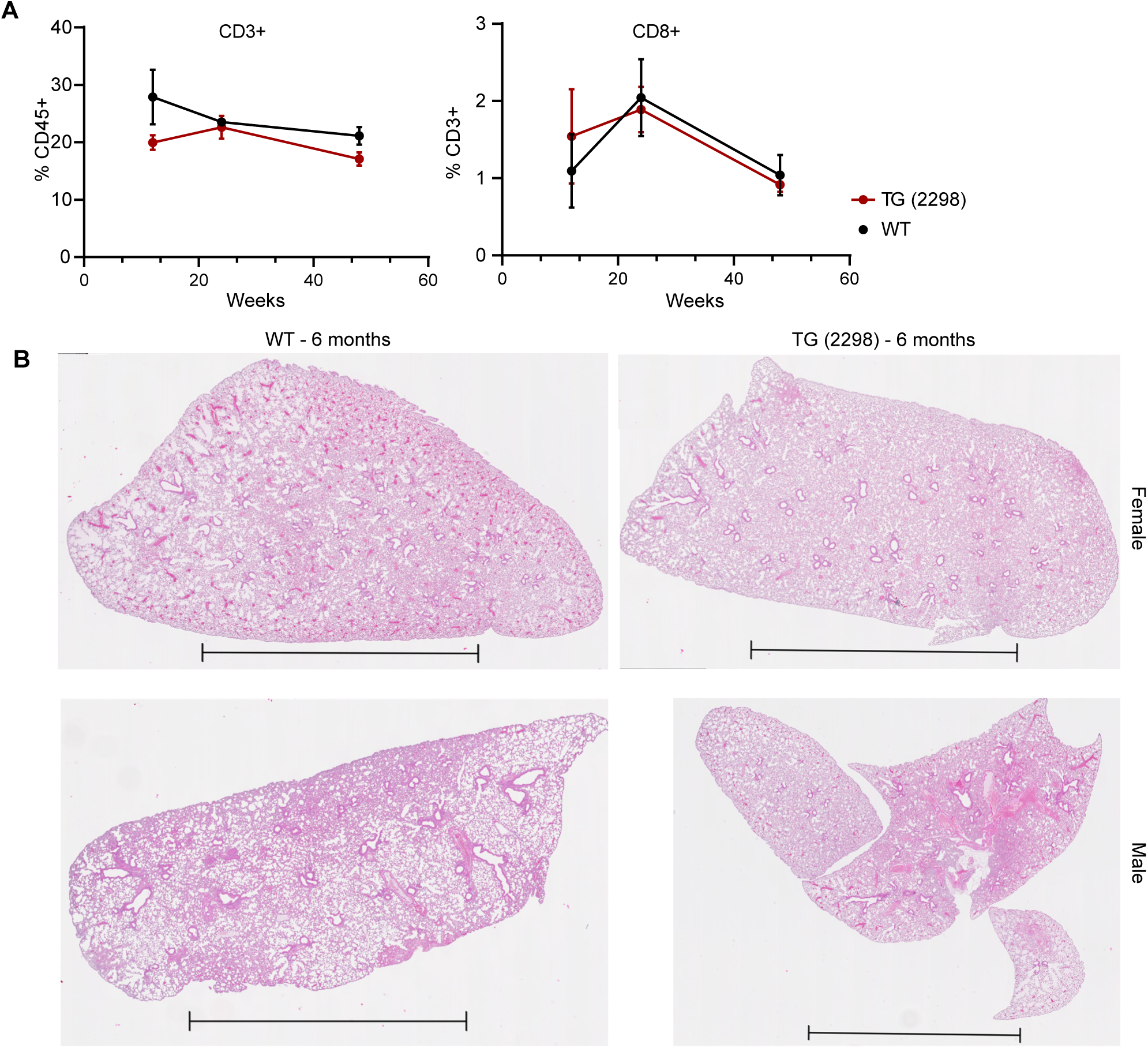
Lungs from TAM-treated TG mice show no inflammation over time at steady states. A) Percentage of CD3+ (gated from CD45+ cells) and CD8+ (gated from CD3+ cells) T cells in the lungs of TG and WT mice at different ages after TAM treatment (n=3 for 12 weeks; n=6 for 24- and 48-week time points). B) Hematoxylin-eosin staining of a lung tissue section of 24-week-old TG and WT TAM-treated mice. Scale bar = 5mm.

Our findings from the transgenic mouse model, which overexpresses MHC I in AT2 cells, reveal no significant inflammatory patterns under steady-state conditions for at least one year. Additionally, there were no modifications in LEC fates or proportions. These results suggest that MHC I overexpression in AT2 cells does not adversely impact lung homeostasis under normal conditions, providing valuable insights into the role of MHC I in lung biology, at least in the clean environment of an academic animal core facility.

## Discussion

We created a transgenic mouse model that overexpresses surface MHC I thanks to the upregulation of NLRC5. While the role of NLRC5 in MHC I regulation has been extensively explored in hematolymphoid cells (Biswas et al., 2012; Ludigs et al., 2015; Meissner et al., 2010), our study likely provides the first evidence of NLRC5’s role in basal MHC I regulation in LECs under steady-state conditions. By breeding our *LSL-Nlrc5* mice with *Sftpc-CreER* mice, we achieved an approximately 10-fold increase in MHC I at the surface of AT2. Genes crucial for the MHC I presentation machinery (*B2m*, *Psmb8*, *Psmb9*, *Tap1*) were significantly upregulated. Our approach offers a versatile model for modulating MHC I levels across various cell types using Cre recombinase expression. Incorporating the mCherry reporter allows rapid validation of transgenic NLRC5 expression, even in tissues already expressing this protein. Our study demonstrates that MHC I overexpression in AT2 cells is both rapid and stable in our transgenic model, enabling long-term investigations into the effects of MHC I overexpression. The duration of transgene expression is influenced by the self-renewing capacity of each cell type, which is notably prolonged in AT2 cells (Barkauskas et al., 2013). Consequently, the stability of transgene expression may vary across different cell types.

Previous studies have shown that LECs express low levels of MHC I, significantly increasing during inflammation and chronic respiratory diseases (Mathé et al., 2022, 2024). While these studies provide new insights into the mechanisms regulating MHC I in LECs, a fundamental question remains: Is MHC I overexpression an initiator or a consequence of the disease? Our transgenic mouse model shows that increasing MHC I does not induce significant pathology or inflammation under steady-state conditions. This finding challenges the notion that MHC I is a primary initiator of chronic respiratory diseases. However, these pathologies predominantly affect older subjects after repeated lung injuries from factors such as tobacco smoke and viral exposure (Ishikawa et al., 2021; Linden et al., 2019; Zuo et al., 2014). Future studies should investigate the effects of these environmental factors on transgenic mice compared to WT counterparts to evaluate the role of MHC I overexpression in disease progression and severity. Furthermore, simultaneously upregulating MHC I in various LEC types, not just AT2 cells, may provide additional insights since multiple LEC types increase MHC I in chronic respiratory diseases. This simultaneous overexpression is achievable by crossing *Sftpc-CreER* mice with *Scgb1a1-Cre* mice, as demonstrated in prior research (Barkauskas et al., 2013; Perl et al., 2009). Our model also offers numerous perspectives to investigate the role of MHC I overexpression in other diseases where MHC I is overexpressed, such as inflammatory bowel diseases, myopathies, or type I diabetes, by increasing MHC I in different cell types outside the lungs.

Given that MHC I downregulation is a common mechanism for cancer and viral immune evasion (Cornel et al., 2020; Koutsakos et al., 2019; Lee et al., 2020), and since the lung epithelium (which typically exhibits low MHC I levels) is particularly susceptible to these diseases, our transgenic mouse model could provide valuable insights into the impact of MHC I levels on cancer and infection susceptibility and severity. Cancer immunotherapies target regulation of MHC I expression, but while increasing MHC I stimulates CD8+ T cells response, it can also break tissue tolerance to self-antigens, especially in tissue where MHC I is initially low (Bassani-Sternberg et al., 2015; Caron et al., 2011; Cebrián et al., 2014; Stern et al., 2024). This study shows a therapeutic window of approximately 10-fold to increase MHC I in AT2 cells without inducing pathology. Such a model of mice overexpressing MHC I in specific tissues could be particularly relevant in optimizing therapies that affect MHC I and minimizing adverse immune effects in cancer treatments (Garrido et al., 2016). This work could pave the way for new, more personalized diagnostic and predictive strategies.

Overall, we have created a new and robust model of in vivo MHC I surface overexpression, which is cell-specific and stable over time. Our analysis of MHC I increase in AT2 cells demonstrated two key points: (i) NLRC5 is a crucial regulator of MHC I and associated genes in LECs, and (ii) elevated MHC I levels in AT2 cells did not induce pathology or inflammation under steady-state conditions. This model offers many perspectives to understand better the role of MHC I expression in immunopathologies, tissue tolerance, and therapeutic development.

## Competing Interests and Funding

Authors declare no competing interests. This work was supported by grant FDN-148400 from the Canadian Institutes of Health Research (to CP).

JM, SB, and CP conceptualized and designed the study. JM conducted experiments, analyzed results, created figures, and wrote the initial draft of the manuscript. JM, SB, and MSL design the vector. MSL performed primer design and contributed to result discussions. CC conducted mice genotyping and PCR analyses. AK helped establish flow cytometry protocols. SH performed microinjections in mouse embryos. CP and SB provided overall supervision for the study. All authors critically reviewed and approved the manuscript.

## Acknowledgments

We thank the dedicated staff at the IRIC core facilities, including the genomics, flow cytometry, microscopy, and histology research units, for their invaluable assistance and insightful discussions. We thank the IRIC animal core facility team for their unwavering support throughout the project.

## Notes

### Competing Interest Statement

The authors have declared no competing interest.

