## Supplementary material for "Transgenic Inducible MHC I Overexpression in Mouse Alveolar Type 2 Cells"

**Running title: Mice Overexpressing MHC I**

Justine Mathé*†‡, Sylvie Brochu*†, Marc K. Saba-El-Leil*†, Caroline Coté*†, Amrita Karia*†, Sébastien Harton*† and Claude Perreault*†.

*Institute for Research in Immunology and Cancer (IRIC), Université de Montréal, Montréal, Quebec, Canada.

†Department of Medicine, Université de Montréal, Montréal, Quebec, Canada

‡Corresponding author:

Justine Mathé : T +438 530 5226,

Sylvie Brochu : T + 514 343 6111 local 0594,

ORCIDs: 0000-0002-4482-1148 (J.M.); 0000-0002-4665-0936 (S.B.); 0000-0001-9453-7383 (C.P.)

Competing Interests and Funding: Authors declare no competing interests. This work was supported by grant FDN-148400 from the Canadian Institutes of Health Research (to CP).

**Supplemental information**

| Founder | % of TG mice | Sexe |
|---|---|---|
| 2298 | 50 | Female |
| 2283 | 44.4 | Female |
| 2287 | 51.8 | Male |
| 2291 | 68.75 | Male |
| 2293 | 15 | Male |

**Table S1. Transmission of the transgene.** Percentage of mice expressing the transgene in the progeny from each founder (F1).

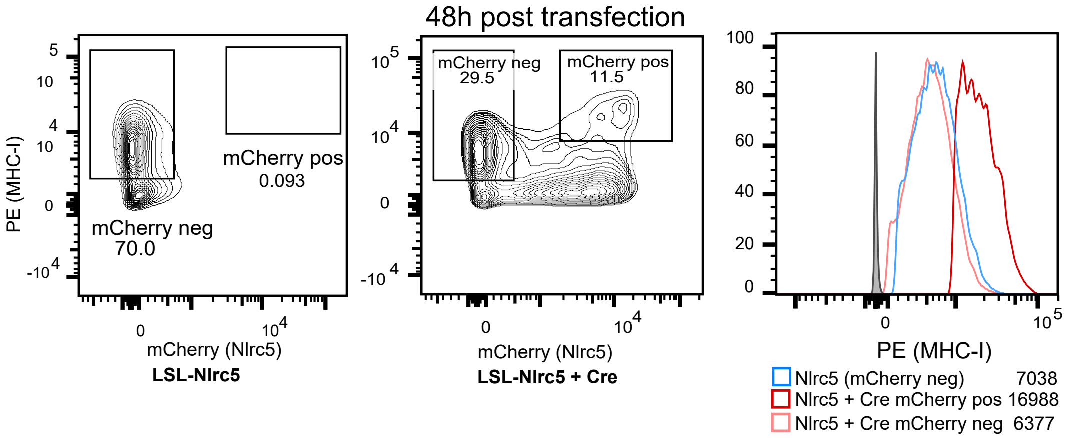


**Figure S1. In vitro validation of the construct in HEK-293 cells.** Percentage of MHC I+ mCherry+ HEK-293 cells transfected with the *LSL-Nlrc5-mCherry* vector with or without the Cre enzyme 48 hours post-transfection.
